# Human Gut Microbiota of Healthy Individuals Show Resilience to Acute Aflatoxin B1 Exposure Despite Solvent-Driven Community Shifts

**DOI:** 10.64898/2026.10.26.760590

**Authors:** Hannah Glesener, Nicole Dorn, Qiwen Cheng, Lee E. Voth-Gaeddert, Rosa Krajmalnik-Brown

## Abstract

Aflatoxin B1 (AFB1) is a potent mycotoxin linked to severe systemic health issues and gut microbiome disruption in in vivo models. However, it remains unclear whether AFB1 directly impairs human gut microbial composition and function or if observed dysbiosis is host-mediated. The objective of this work was to investigate the direct impact of acute AFB1 exposure on human gut microbiota from healthy individuals using anaerobic batch bioreactors. Fecal inoculum from 17 healthy adults was exposed to AFB1 concentrations ranging from 1,000 to 5,000 ppb, delivered in two solvents (ethanol and corn oil) under conditions simulating the large intestine. Metabolic activity was assessed via short-chain fatty acid (SCFA) and gas production measurements, while community structure was assessed using 16S rRNA gene amplicon sequencing. AFB1 concentrations remained stable throughout the incubations, indicating no significant microbial degradation of AFB1. Results showed that AFB1 did not significantly alter microbial SCFA output or community composition compared to solvent-only controls. Observed shifts in microbial profiles were primarily driven by the solvents used to bring AFB1 into solution rather than by the toxin. These findings demonstrate that healthy human gut microbial communities are resilient to direct, acute AFB1 exposure even at concentrations exceeding typical dietary levels. This suggests that microbiome disruption reported in vivo may be an indirect consequence of host-mediated factors, such as reactive metabolites or systemic immune responses, rather than direct toxicity by the parent compound.

**Importance:** Aflatoxin B1 is a major global health threat, particularly in regions with high dietary exposure, where it is associated with liver cancer and childhood growth stunting. While research has linked aflatoxin exposure to changes in the gut microbiome, the mechanisms driving this “dysbiosis” are not known. This work provides evidence that the parent aflatoxin molecule does not directly affect healthy human gut microbes, even at very high concentrations. The observed resilience of these microbial communities suggests that the damage reported in living organisms is likely caused by host-specific responses to the toxin or by toxic byproducts created during liver detoxification. Furthermore, microbially driven detoxification of the parent aflatoxin molecule was not observed. Distinguishing between direct and indirect toxicity is essential for developing effective strategies and medical interventions to mitigate the health impacts of aflatoxin exposure worldwide.

## Introduction

Aflatoxin B1 (AFB1) is a potent globally prevalent mycotoxin produced by *Aspergillus flavus* and *A. parasiticus* that commonly contaminates staple foods such as maize, wheat, and nuts (1–6). Chronic dietary exposure to AFB1 is a major public health concern in tropical and subtropical regions and has been strongly linked to hepatocellular carcinoma, immunosuppression, and childhood growth stunting (7–9). Increasingly, AFB1 is also implicated in alterations to the gut microbiome, which may further exacerbate its systemic toxicity by impairing intestinal barrier function and nutrient absorption (10–12). Growth stunting, one of the surmised consequences of early-life AFB1 exposure, is associated with long-term cognitive deficits, reduced productivity, and increased risk of chronic disease, making it a critical developmental and global health issue (13, 14). Despite extensive studies of AFB1’s hepatotoxicity, the direct mechanisms by which it impacts human gut microbiota remain poorly understood.

Several in vitro studies have examined how individual microbial isolates respond to AFB1 exposure. Many strains, particularly among lactic acid bacteria such as *Lactobacillus plantarum*, *Bifidobacterium* spp., and *Pediococcus pentosaceus*, have demonstrated the ability to bind AFB1 through surface adsorption (e.g., non-covalent binding to peptidoglycan) without showing signs of inhibited growth or toxicity, even at concentrations up to 5 µg/mL (5,000 ppb) (15, 16). Additionally, some microbes can detoxify AFB1. For instance, *Bacillus amyloliquefaciens* (from fermented pickles) and *Microbacterium proteolyticum* (from bovine feces) degrade >75% of AFB1 within 24–72 hours, converting it to nontoxic metabolites (17, 18). Studies using microbial consortia from compost have shown high tolerance for and effective degradation of AFB1, with minimal community composition disturbance (19, 20).

In contrast, in vivo animal studies have consistently shown that AFB1 exposure leads to gut microbiota disruption. In F344 rats and piglets, low-grade AFB1 exposure (0.0075-1 µg/mL; 75-1,000 ppb) decreases microbial diversity, depletes beneficial taxa like *Lactobacillus*, and reduces short-chain fatty acid (SCFA) production (21, 22). These disruptions often correlate with intestinal inflammation, increased gut permeability, and impaired bile acid metabolism. In mouse models, AFB1 alters the composition and function of the gut microbiota and exacerbates host metabolic dysfunction, but these effects often rely on host signaling pathways, such as farnesoid X receptor suppression or cytochrome P450 enzyme activation (23). Importantly, such host-dependent effects make it difficult to disentangle whether AFB1 exerts direct toxicity on gut microbes or whether microbiome changes are downstream consequences of host processing and immune responses.

This contrast between the resilience of in vitro microbes and the vulnerability of in vivo microbes to AFB1 exposure highlights a critical gap in our understanding of whether the gut microbiota is intrinsically sensitive to AFB1 or whether the observed disruptions are entirely dependent on host processing (e.g., metabolic and immune modulation). Understanding whether the gut microbiota is directly affected by AFB1 has important implications for food safety, therapeutic design, and environmental toxin risk modeling. In this study, we addressed this gap by testing the hypothesis that direct exposure to AFB1 impairs the composition and metabolic activity of the human gut microbiota. Using anaerobic batch bioreactors inoculated with fecal samples from healthy adults, we exposed the microbial consortia to varying concentrations of AFB1 (up to 5,000 ppb) dissolved in solvent across a 36–48-hour incubation in gut media. To distinguish effects of the parent toxin from those of the solvent vehicle, we evaluated AFB1 using two biologically relevant solvents, ethanol and corn oil (24). We monitored microbial metabolic activity using SCFA production, gas pressure and composition, pH, and quantified AFB1 recovery to assess degradation. Our aim was to determine whether the parent compound AFB1 directly alters microbial composition and metabolism or whether the healthy adult gut microbiota is functionally resilient under acute toxin exposure in vitro.

## Materials and Methods

### Fecal sample collection and preparation

Samples were collected following ethical approval from the Arizona State University (ASU) Institutional Review Board (STUDY00017164). Participants were recruited from ASU and the surrounding community using word-of-mouth, ASU’s “Research Plus Me” platform, and flyers linked to a QR-code eligibility screening form. Inclusion criteria included: age between 18–65 years, average weight (BMI 18.5–30), no gastrointestinal diseases (e.g., diarrhea, constipation, Crohn’s), no medical treatment in the prior two months, and no use of antibiotics or antifungal medications in the previous three months. Exclusion criteria included: pregnancy, children (<18), prisoners, individuals unable to consent, or those with autoimmune, hepatitis, or mental/physical disorders. Once eligibility was confirmed and written informed consent obtained, participants completed intake questionnaires and received stool collection kits along with instructions for sample return within four hours of defecation. Samples were stored at 4 °C temporarily.

Within 12 hours of sample receipt, stool was transferred into an anaerobic glovebox for processing. For each participant, we prepared 10 raw aliquots (1 gram each in 2 mL cryotubes), 10 glycerol-preserved aliquots (1–2 grams in 5 mL of 10% filter-sterilized glycerol), and stored the remaining material as raw bulk in 50 mL Falcon tubes. All aliquots were immediately stored at –80 °C. After collecting a sufficient number of samples, we prepared composite inoculum by pooling one glycerol-preserved aliquot from each of 17 participants in the anaerobic glovebox. From each pooled mixture, 1 mL was aliquoted into 2 mL cryotubes and stored at –80 °C. These composite mixtures were later thawed and used to inoculate bioreactors.

### Bioreactor Inoculation and Setup

A medium designed to simulate the large intestine, adapted from McDonald et al. (2013) (25), was used for all experiments. The medium components were diluted by 1/20^th^ the original specifications because the formulation was originally designed for continuous, not batch reactors. For each AFB1 experiment, the preparation and sampling followed a generic processing timeline. Reactors were inoculated with slurry prepared from aliquots of the 17 participants’ fecal samples that were preserved in 10% glycerol, thawed on ice and mixed 1:1 by volume with media. The fecal slurry was incubated for 24 hours at 37 °C in the dark with agitation to promote microbial acclimation. Following incubation, 1.2 mL of slurry was inoculated into each bioreactor vessel inside an anaerobic glovebox. Bioreactors were run for 36-48 hours to simulate a typical gut transit time.

Crystalline solid AFB1 (Item No. 11293, Cayman Chemical, Ann Arbor, MI, USA) was kept in its original container (1 mg in 4 mL amber vial) and either ≥ 99.50 % ethanol (CAS: 64-17-5, Thermo Fisher Scientific, Pittsburgh, PA, USA) or corn oil, NF (CAS: 8001-30-7, Spectrum, Stamford, CT, USA) solvents were dispensed into an AFB1 container in an anaerobic glovebox to achieve a concentrated solution of 250,000 ppb. The stock was dispensed into a serum bottle with 60 mL of media to achieve the desired concentration, for example, 240 µL for a 1,000 ppb dosage.

### Sampling Procedures

Pressure was read with a barometer (General, Cincinnati, OH, USA). Gas was measured on a gas chromatograph (Agilent, Santa Clara, CA, USA) using 200 µL of sample headspace. Liquid samples were immediately processed for downstream analysis by expelling 1 mL of sample into 2 mL microcentrifuge tubes. A combination semi-micro pH electrode on a benchtop meter (Orion Star A211, Thermo Scientific, Waltham, MA, USA) was used to measure pH. Samples were centrifuged at 10,000 rpm for 10 min, then sterile-filtered using 0.22 µm PTFE filters attached to a sterile 1 mL syringe, and transferred into a 2 mL amber auto-sampler vial for SCFA analysis. For aflatoxin analysis, 150 µL of the samples were withdrawn with a sterile pipette and injected into a second amber autosampler vial containing a 150 µL spring vial insert. The remaining biomass was immediately set on ice, then stored at −80 °C for DNA extractions. Samples were taken every 12 to 24 hours.

### Chemical Analyses

Gas composition was measured on an Agilent Gas Chromatograph (8890 GC System, Agilent Technologies, Santa Clara, CA, USA) configured with FID and TCD detectors and argon as a carrier gas via manual injection of 200 µL gas sample. Gas flow rate was 7 mL/min at 280 °C over a 14.833 min run time. Standard curves for TCD detected gas (H_2_) and FID detected gas (CH_4_ and CO_2_) were calculated and had R^2^ > 0.95.

Aflatoxin and SCFAs were measured on an Agilent High Performance Liquid Chromatograph (1260 Infinity II HPLC) configured with a refractive index detector and diode array detector (DAD). Aflatoxin was measured on an Agilent Eclipse XDB-C18 4.6mm × 150mm, 3.5 μm Rapid Resolution column, using the DAD, according to previously published methods (26). A certified reference material (CRM46304; Sigma-Aldrich, St. Louis, MO, USA) was used to calculate a standard curve for AFB1 with R^2^ > 0.95.

SCFAs were measured on the HPLC with adaptations from previously published methods (27, 28). The system was equipped with a Fast Acid Analysis HPLC column, 100 x 7.8 mm (Bio-Rad Laboratories, Hercules, CA, USA). An autosampler set at 8 °C injected 10 µL of sample into a mobile phase of 0.005 mM H_2_SO_4_ at a 0.5 mL/min flow rate for 35 min. SCFAs were detected with the DAD detector set at 210 nm with a 360 nm reference wavelength. A certified reference material (CRM46875; Sigma-Aldrich, St. Louis, MO, USA) was used to calculate a standard curve with R^2^ > 0.95 for acetic acid, propionic acid, and butyric acid (referred to in this manuscript as acetate, propionate and butyrate).

### Microbiome Sequencing and Analysis

Microbial DNA was extracted from biomass obtained from 1 mL liquid samples using the DNeasy PowerSoil Kit (QIAGEN, Hilden, Germany) according to the manufacturer’s recommendations as previously described (29). Full-length 16S rRNA gene amplicon libraries were prepared using the PacBio Kinnex 16S kit following the manufacturer’s protocol using barcoded 27F/1492R Kinnex primers. Amplicons were pooled and converted into SMRTbell libraries through successive adapter ligation, cleanup, and array formation steps, including Kinnex array assembly to concatenate 16S rRNA gene amplicons into linear molecules. Libraries were purified using AMPure PB beads and treated with nuclease according to PacBio guidelines. Final SMRTbell libraries were quantified using a Qubit fluorometer (Thermo Fisher Scientific, Waltham, MA, USA) and assessed for size distribution and quality using a Femto Pulse system (Agilent, Santa Clara, CA, USA). Barcoded libraries were pooled equimolarly and sequenced on a PacBio Revio SMRT cell. Sequencing quality was assessed using FastQC, and summary reports were generated with MultiQC to evaluate metrics including sequence quality scores, duplication levels, and adapter content.

Raw fastq files were processed through PacBio’s HiFi Nextflow pipeline (https://github.com/PacificBiosciences/HiFi-16S-workflow; v0.9) using default parameters. This pipeline used seqkit (2.8.2) and csvtk (0.27.0) for initial FASTQ inspection and metadata handling. Primer trimming and orientation were performed with cutadapt (4.9). Reads were denoised into amplicon sequence variants using DADA2 (1.30.0) within QIIME 2 (2024.10). Low-abundance features were removed using the workflow’s default thresholds, which retain amplicon sequence variants supported by at least five reads across at least one sample.

Taxonomic assignment was performed using the workflow’s Naïve-Bayes “best taxonomy” approach, which integrates assignments from Greengenes2, GTDB r207, and SILVA 138.1 in that order. All downstream processing, visualization, and statistical analyses were performed in R (version 4.5.1). ASV counts, taxonomic assignments, and sample metadata were integrated into phyloseq objects with phyloseq (1.52.0), which was used for alpha and beta diversity analysis. Data wrangling and plotting were conducted with tidyverse (2.0.0).

### Statistical Analyses

All statistics were calculated in R (version 4.5.1) using base R functions and the rstatix package (v0.7.2) for non-parametric post-hoc testing. At each timepoint, data were assessed for normality using the Shapiro-Wilk test on residuals from a one-way ANOVA model. When the Shapiro– Wilk test did not detect a departure from normality (p > 0.05), treatment means were compared using one-way ANOVA with Tukey-adjusted pairwise comparisons, otherwise a Kruskal-Wallis test was used with Dunn’s test and Bonferroni correction. For microbiome statistics, Bray–Curtis and Jaccard distance calculations, ordinations, and PERMANOVA tests were performed with vegan (2.7.2) as previously described (30).

## Results and Discussion

### AFB1 did not significantly change SCFA production by gut microbiota

Because aflatoxin B1 (AFB1) has limited solubility in water, we tested two physiologically relevant solvents, corn oil and ethanol, to dissolve AFB1 and deliver the calculated toxin dose to gut microbiota in vitro. As seen in Figure 1a, overall, direct exposure to AFB1 in solvent did not significantly alter the production of the primary SCFAs (acetate, propionate, and butyrate) when compared to the solvent-only condition (p > 0.05). One way ANOVA revealed significant differences for some timepoints and acid concentrations when comparing all conditions (e.g., acetate at 24h, p=0.0083; 36h, p=0.0047), but follow-up pairwise comparison testing revealed that no significant differences in SCFA production existed between the solvent conditions and their AFB1-treated counterparts (e.g., corn oil vs. corn oil + AFB1) at any timepoints (Fig. 1a, SI Fig. 1). Instead, the differences in SCFAs measured were driven by the presence of solvents. Unlike SCFAs, gas production did not differ significantly between any conditions (Fig. 1b).

**Figure 1.**
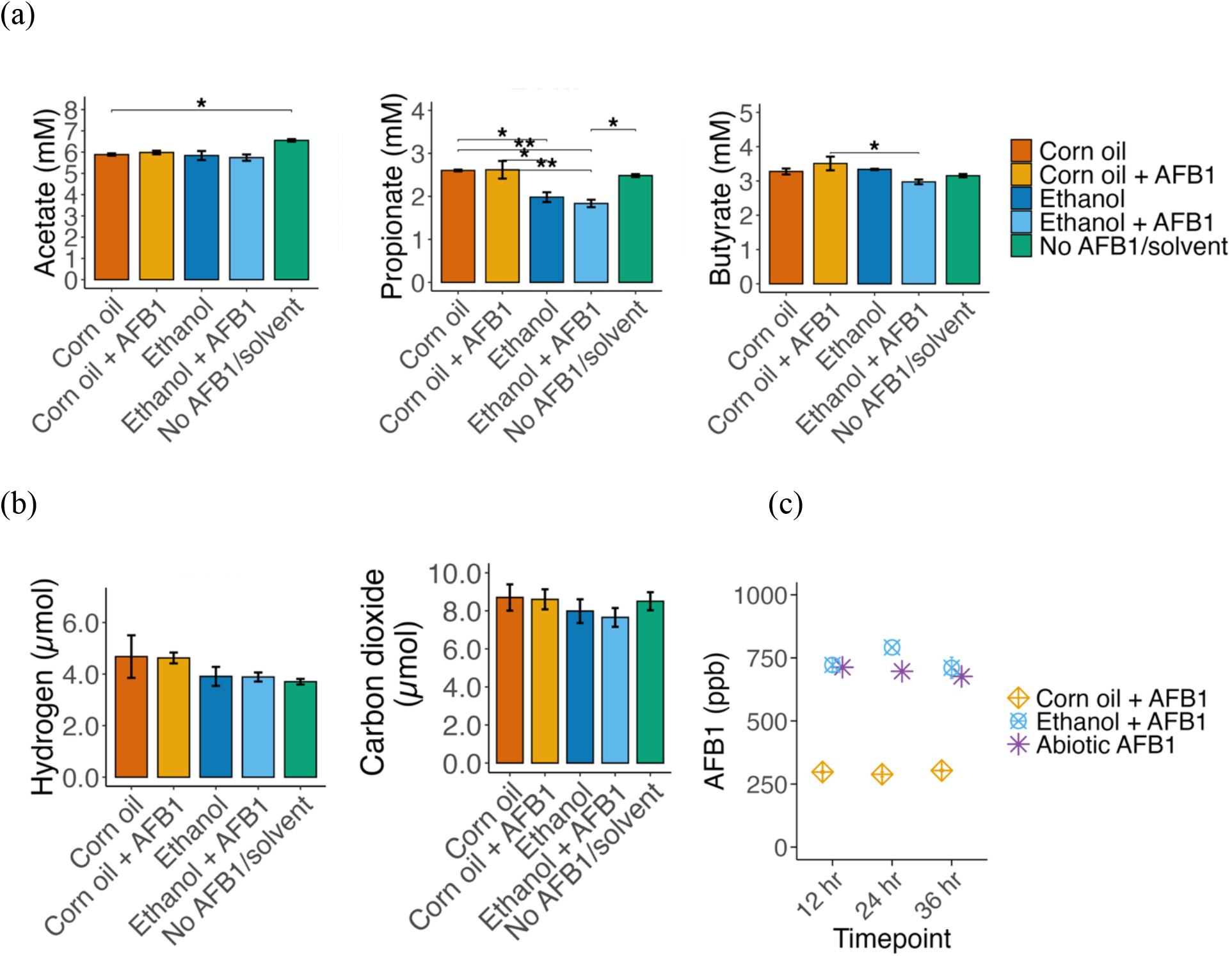
Metabolic activity in response to AFB1 exposure in complex gut medium. (a) Acetate, propionate, and butyrate concentrations measured at 24 hours in bioreactors containing corn oil or ethanol with 1,000 ppb AFB1, solvent controls, and an untreated control of media and microbes with no solvent or toxin. All reactors were run in triplicate. (b) Hydrogen and carbon dioxide amounts at 24 hours across treatment groups. (c) Measured AFB1 concentrations across experimental timepoints. Data are presented as mean ± standard error of the mean (SEM). Statistical significance was assessed using one-way ANOVA with Tukey’s HSD post-hoc test or Kruskal–Wallis test with Dunn’s post-hoc test and Bonferroni correction, as appropriate. Asterisks indicate significant pairwise differences: *p < 0.05, **p < 0.01, ***p < 0.001.

A key observation across all metabolic measurements was the consistency in functional output, measured by SCFAs and gas production over time. The SCFAs and gas production trajectories for all treatment groups followed a similar pattern, with no condition exhibiting a strong deviation or detrimental effect (e.g., no production or lack of increase over time). For instance, butyrate concentration measured at 24 hours in the condition without AFB1 or solvent was 3.15 ± 0.05, the corn oil + AFB1 was 3.5 ± 0.2 and ethanol + AFB1 was 2.97 ± 0.06. This implies that AFB1 does not have a detrimental effect on SCFA production within the tested hydraulic retention times and that the community involved in this process is functionally resilient to this exposure. It is also important to note that AFB1 (dissolved in solvent) remained stable throughout the experiment, measured in microbially active reactors and an abiotic control, indicating limited to no microbial degradation of the parent toxin (Fig. 1c).

This functional resilience is consistent with previous findings from environmentally derived microbial consortia, which tolerated high concentrations of AFB1 with minimal community disturbance (19, 20). In the present study, the solvent vehicle, rather than the AFB1 exposure, was the primary driver of metabolic variation.

### AFB1 did not significantly change microbial community composition

We assessed whether AFB1 exposure altered gut microbial community composition. Beta diversity analyses indicated that microbial communities clustered primarily by solvent type rather than AFB1 treatment. When all conditions were analyzed together, community composition separated strongly by solvent background (Bray–Curtis R² = 0.683, p = 0.001; Fig. 2a). In contrast, within-solvent comparisons revealed no significant effect of AFB1 treatment on overall community structure in either the corn oil (R² = 0.063, p = 0.322; Fig. 2b) or ethanol (R² = 0.098, p = 0.165; Fig. 2c) conditions. Similar patterns were observed using Jaccard distance, indicating that the observed community structure was not driven solely by changes in dominant taxa (SI Fig. 2a).

**Figure 2.**
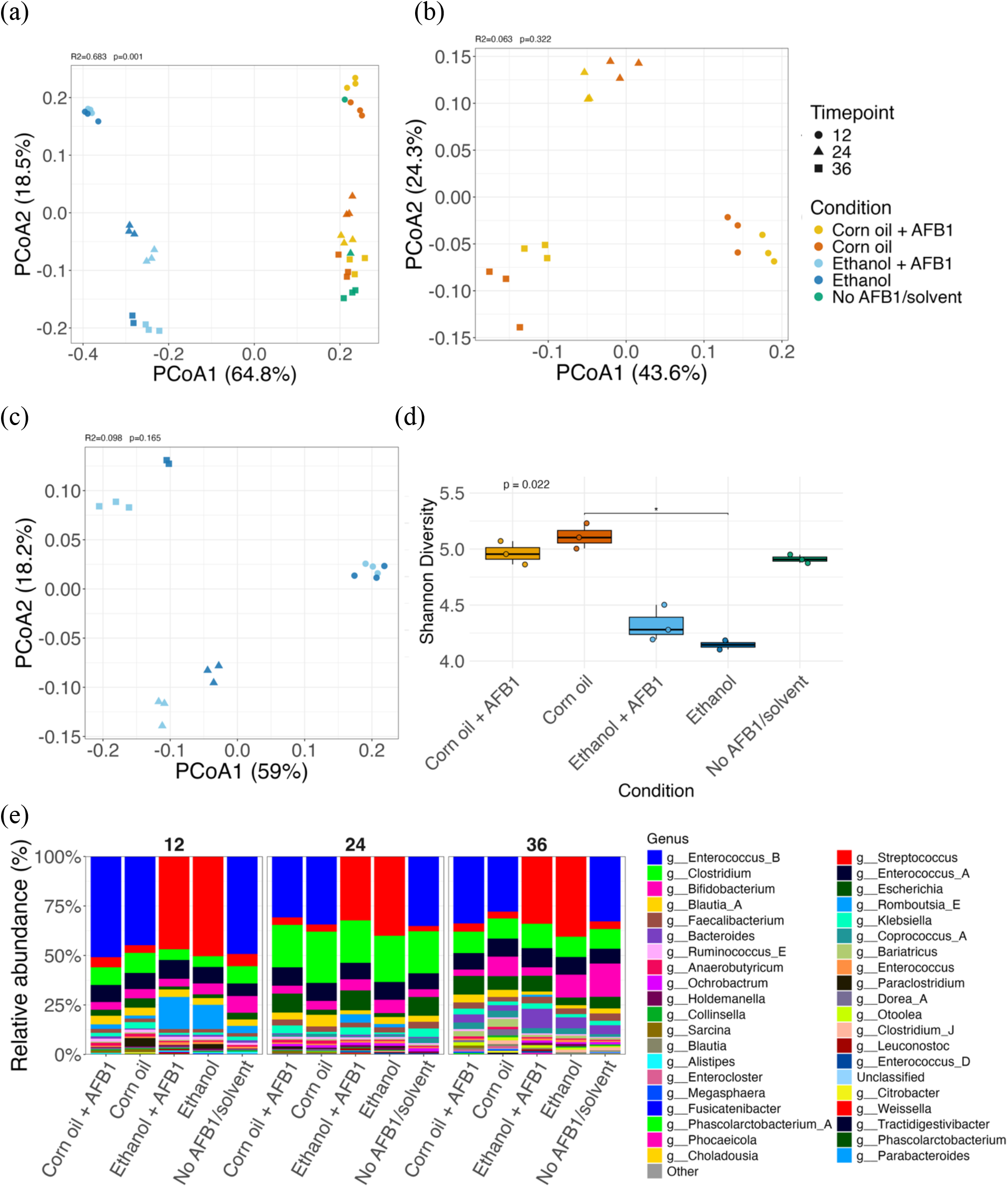
Microbial community composition and diversity across solvent and AFB1 treatments and an untreated control of media and microbes with no solvent or toxin. (a) Principal coordinates analysis (PCoA) of Bray–Curtis dissimilarity including all treatment groups, (b) corn oil and (c) ethanol solvent conditions. Differences in community composition were assessed by PERMANOVA. (d) Shannon diversity index at 36 hours. Differences among conditions were assessed using the Kruskal–Wallis test. (e) Relative abundance of the top 20 genera across solvent conditions ± 1,000 ppb AFB1 over time. Asterisks indicate significant pairwise differences: *p < 0.05, **p < 0.01, ***p < 0.001.

Consistent with these findings, relative abundance profiles of the top 20 genera clustered primarily by solvent type rather than by AFB1 treatment (Fig. 2e). Within each solvent group, AFB1-treated communities closely resembled their respective solvent-only controls across all timepoints, with no clear blooms or losses of dominant taxa. Alpha diversity metrics differed among experimental groups at 36 hours (e.g., Shannon diversity at 36 h, p = 0.022; Fig. 2d).

However, these differences were attributable to solvent background rather than AFB1 exposure. Within each solvent, AFB1-treated samples did not show consistent reductions in richness, diversity, or evenness compared with their matched controls (SI Fig. 3). Together, these findings indicate that solvent context was the primary driver of microbial community structure, whereas short-term AFB1 exposure did not induce notable taxonomic restructuring, consistent with the stability observed in metabolic output.

### A five-fold increase in AFB1 dose failed to significantly inhibit microbial activity

To investigate whether the lack of microbial response to AFB1 was due to an insufficient toxin concentration, we performed a subsequent experiment increasing the AFB1 concentration five-fold to 5,000 ppb in ethanol. Despite this substantial increase in AFB1 concentration, microbial metabolic output remained robust, demonstrating resilience to high-dose AFB1 exposure (Fig. 3).

**Figure 3.**
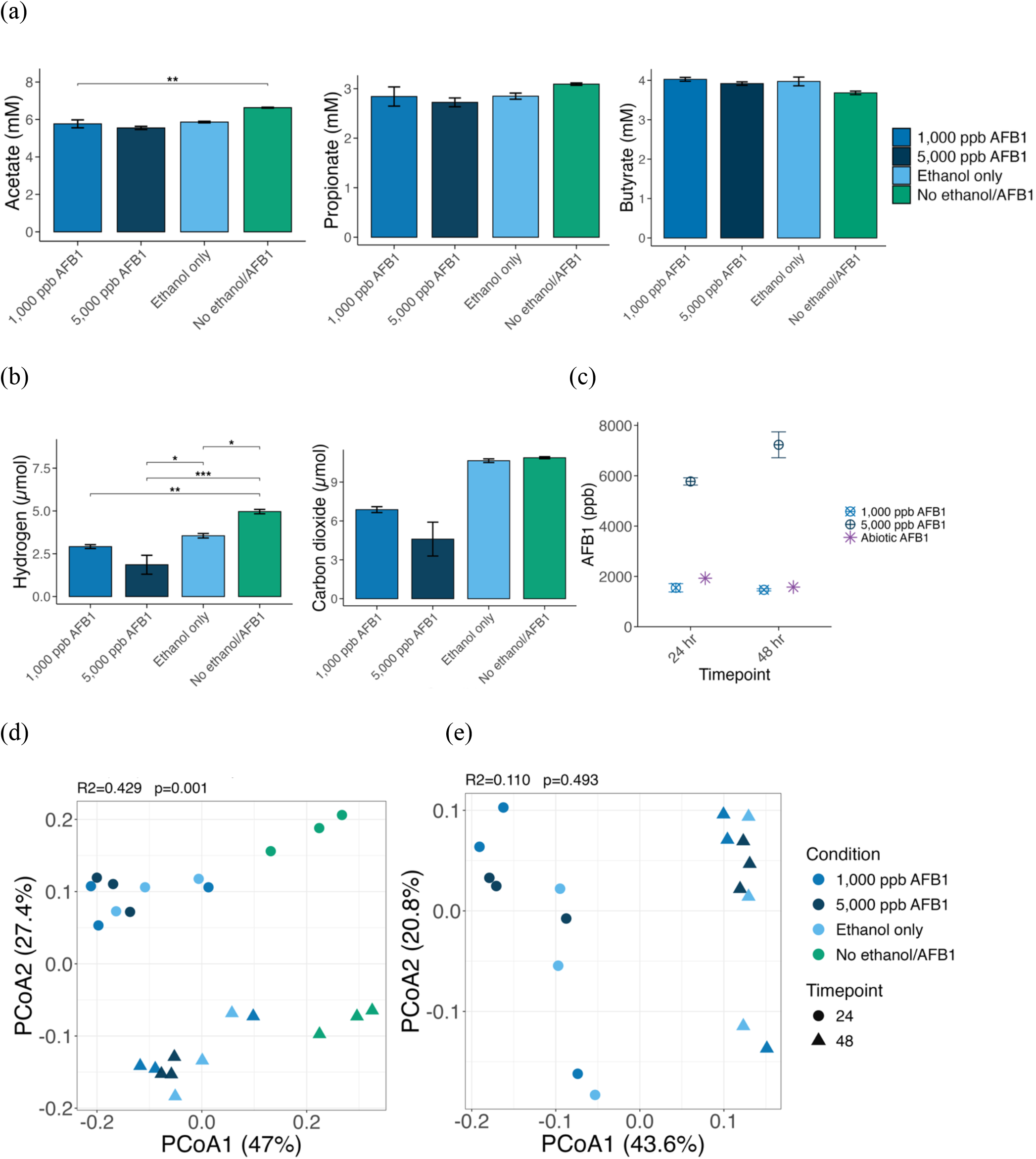
Metabolic activity and microbial composition changes in response to AFB1 exposure in complex gut medium. Bioreactors treated with 1,000 ppb AFB1 in ethanol, 5,000 ppb AFB1 in ethanol, an ethanol only control, and an untreated control with no ethanol or AFB1. All reactors were run in triplicate. (a) Acetate, propionate, and butyrate concentrations at 24 hours and (b) Hydrogen and carbon dioxide amounts at 24 hours. Error bars represent mean ± SEM. Statistical significance was assessed using one-way ANOVA with Tukey’s HSD post-hoc test or Kruskal-Wallis test with Dunn’s post-hoc test and Bonferroni correction, as appropriate. (c) AFB1 concentration at all timepoints. (d-e) Bray-Curtis beta diversity PCoA plots. Differences in community composition were assessed by PERMANOVA. Asterisks indicate significant pairwise differences: *p < 0.05, **p < 0.01, ***p < 0.001.

Across all active bioreactors, SCFA and gas production increased between 0 and 24 hours and remained stable from 24 to 48 hours, with no clear signs of inhibition in the AFB1-treated groups (Fig. 3a, 3b and SI Fig. 4). There were no statistically significant differences in propionate or butyrate production when comparing the 1,000 ppb and 5,000 ppb AFB1 conditions at any timepoint. A minor but statistically significant decrease in acetate was observed in the 5,000 ppb group (6.07 mM) compared to the 1,000 ppb group (7.56 mM) at 48 hours (Tukey HSD, p < 0.001), but this did not reflect a broader pattern of metabolic inhibition (SI Fig. 4). Similarly, no major differences were found in gas production (carbon dioxide and hydrogen) between the two doses.

Although most comparisons were not statistically significant, a few measurements were numerically lower in the 5,000 ppb treatment. For instance, at 24 hours, the mean acetate concentration in the reactor with no solvent or AFB1 was 6.63 mM, compared to 5.86 mM in the ethanol only condition, 5.76 mM in the 1,000 ppb group, and 5.55 mM in the 5,000 ppb group (Fig. 3a). A similar pattern was observed for hydrogen and carbon dioxide amounts at 48 hours, with the 5,000 ppb group measuring 2.94 µmol H_2_ and 8.1 µmol CO_2_, compared to 2.97 µmol H_2_ and 9.86 µmol CO_2_ in the reactor with no solvent or AFB1, 3.80 µmol H_2_ and 10 µmol CO_2_ in the 1,000 ppb group, and 2.96 µmol H_2_ and 10.25 µmol CO_2_ in the ethanol only condition (SI Fig. 4). Despite slightly lower measurements in some conditions, the bioreactors remained highly active under all conditions, continuing to produce substantial quantities of SCFAs and gases.

Consistent with the metabolic measurements, beta diversity analyses revealed no significant effect of increasing AFB1 concentration on overall microbial community structure. When all conditions were analyzed together, community composition differed significantly (Bray–Curtis R² = 0.429, p = 0.001; Fig. 3d), reflecting separation of the untreated control from the solvent-containing treatments. Similar results were obtained using Jaccard distance (R² = 0.359, p = 0.001; SI Fig. 2), indicating that these differences were not driven solely by changes in dominant taxa. However, when the analysis was restricted to solvent-containing bioreactors, community structure did not differ significantly among the 1,000 ppb AFB1, 5,000 ppb AFB1, and ethanol only conditions using either Bray–Curtis (R² = 0.110, p = 0.493; Fig. 3e) or Jaccard distance (R² = 0.115, p = 0.412; SI Fig. 2b). Together, these findings indicate that increasing AFB1 concentration five-fold did not induce detectable restructuring of the microbial community, regardless of whether community differences were assessed using abundance-weighted or presence/absence metrics.

As in the previous experiment, AFB1 concentrations remained stable over the 48-hour period, confirming that the absence of a measurable microbial response was not attributable to degradation of the compound (Fig. 3c). Collectively, these results indicate that even at a high concentration, direct exposure to the AFB1 parent molecule did not substantially alter microbial community composition or impair overall metabolic activity in this in vitro model of a healthy human gut microbiota.

### Implications and limitations of direct AFB1 exposure

Our findings suggest that healthy human gut consortia are resilient to acute, direct exposure to unmetabolized AFB1. However, several limitations should be considered. First, this study evaluated short-term exposure within an average gut hydraulic retention time, and longer-term or repeated exposures may yield different microbial responses. Second, the inoculum was derived from healthy adult donors; therefore, these findings cannot be directly extrapolated to vulnerable populations such as infants, malnourished individuals, or individuals with pre-existing dysbiosis. Within these constraints, our results suggest that microbiome alterations reported in vivo may arise as downstream consequences of host detoxification products, recirculated AFB1 metabolites, immune signaling, or altered intestinal physiology rather than from direct exposure to unmetabolized AFB1 itself (31).

This distinction has important implications for therapeutic development. Given the rapid absorption of AFB1 in the small intestine, opportunities for microbiota-directed strategies aimed solely at binding or degrading unmetabolized AFB1 are likely to be limited (32). Instead, future interventions may warrant greater focus on mitigating damage associated with host-derived AFB1 metabolites while supporting recovery of microbial community structure and function through microbiota-targeted nutritional or ecological approaches. In addition, strategies that promote or maintain a resilient gut microbial community before exposure may help preserve microbiome stability and improve host resilience to the downstream consequences of AFB1 exposure, particularly in susceptible populations.

### Conclusion

This study provides evidence that healthy human gut microbial communities are resilient to acute, direct exposure to AFB1 in vitro. Across concentrations up to 5,000 ppb, AFB1 did not substantially alter microbial community composition or SCFA production, while solvent vehicle was a stronger driver of observed variation. AFB1 also remained stable throughout incubation, indicating that microbial resilience was not attributable to degradation of the parent compound. Together, these findings suggest that microbiome alterations reported in vivo may involve host-mediated processes rather than direct toxicity of AFB1 to the gut microbiota.

## Supplemental Material

Supplementary Material for this manuscript is available as a separate file, which includes Supplementary Figures 1 through 4.

## Data Availability

The raw 16S rRNA gene amplicon sequencing data generated in this study have been submitted to the National Center for Biotechnology Information (NCBI) Sequence Read Archive (SRA) database. The BioProject accession number will be added upon completion of NCBI processing (submission ID: SUB16566671).

## Ethics Approval

Human fecal samples were collected following ethical approval from the Arizona State University (ASU) Institutional Review Board (STUDY00017164).

## Author contributions

Conceptualization: H.G., Q.C., R.K.B., and L.V.G. Funding acquisition: Q.C., R.K.B., and L.V.G. Sampling: H.G. and N.D. Methodology: H.G., N.D., Q.C., R.K.B., and L.V.G. Data analysis: H.G. and N.D. Supervision: Q.C., R.K.B., and L.V.G.. Writing original draft: H.G. Manuscript review and editing: H.G., N.D., Q.C., R.K.B., and L.V.G.

## Acknowledgments

The authors would like to thank Dr. Peter Rohloff, Dr. Ann Miller, Dr. Gabriela Montenegro, and Olga Torres for advice in experimental design and for collaborative grant application.

## References

1. Temba BA, Darnell RE, Gichangi A, Lwezaura D, Pardey PG, Harvey JJ, Karanja J, Massomo SMS, Ota N, Wainaina JM, Fletcher MT, Kriticos DJ. 2021. The Influence of Weather on the Occurrence of Aflatoxin B1 in Harvested Maize from Kenya and Tanzania. Foods 10:216.

2. Daou R, Hoteit M, Bookari K, Joubrane K, Khabbaz LR, Ismail A, Maroun RG, Khoury AE. 2023. Public health risk associated with the co-occurrence of aflatoxin B1 and ochratoxin A in spices, herbs, and nuts in Lebanon. Front Public Health 10:1072727.

3. Monger A, Mongar P, Dorji T, Chhetri V. 2024. The occurrence and human health risk assessment of total and aflatoxin B1 in selected food commodities in Bhutan. Sci Rep 14:16258.

4. Calderón R, Palma P, Godoy M, Vidal M, Rivera A. 2023. Co-occurrence and estimation of the risk of total aflatoxins (B1, B2, G1, and G2) and ochratoxin A in agri-food products consumed in Chile. Food Control 146:109493.

5. Leggieri MC, Toscano P, Battilani P. 2021. Predicted Aflatoxin B1 Increase in Europe Due to Climate Change: Actions and Reactions at Global Level. Toxins 13:292.

6. Jallow A, Xie H, Tang X, Qi Z, Li P. 2021. Worldwide aflatoxin contamination of agricultural products and foods: From occurrence to control. Comp Rev Food Sci Food Safe 20:2332–2381.

7. Smith JW, Groopman JD. 2018. Aflatoxins, p. B9780128012383650304. In Reference Module in Biomedical Sciences. Elsevier.

8. Mohsenzadeh MS, Hedayati N, Riahi-Zanjani B, Karimi G. 2016. Immunosuppression following dietary aflatoxin B1 exposure: a review of the existing evidence. Toxin Reviews 35:121–127.

9. Urugo MM, Teka TA, Lema TB, Lusweti JN, Djedjibegovíc J, Lachat C, Tesfamariam K, Mesfin A, Astatkie T, Abdel-Wahhab MA. 2024. Dietary aflatoxins exposure, environmental enteropathy, and their relation with childhood stunting. International Journal of Food Sciences and Nutrition 1–14.

10. Voth-Gaeddert LE, Torres O, Maldonado J, Krajmalnik-Brown R, Rittmann BE, Oerther DB. 2019. Aflatoxin Exposure, Child Stunting, and Dysbiosis in the Intestinal Microbiome Among Children in Guatemala. Environmental Engineering Science 36:958– 968.

11. Ye L, Chen H, Tsim KWK, Shen X, Li X, Li X, Lei H, Liu Y. 2023. Aflatoxin B _1_ Induces Inflammatory Liver Injury via Gut Microbiota in Mice. J Agric Food Chem 71:10787– 10797.

12. Ye L, Chen H, Wang J, Tsim KWK, Wang Y, Shen X, Lei H, Liu Y. 2024. Aflatoxin B1-induced liver pyroptosis is mediated by disturbing the gut microbial metabolites: The roles of pipecolic acid and norepinephrine. Journal of Hazardous Materials 474:134822.

13. Akbar RR, Kartika W, Khairunnisa M. 2023. The Effect of Stunting on Child Growth and Development. Scientificj 2:153–160.

14. De Onis M, Branca F. 2016. Childhood stunting: a global perspective. Maternal & Child Nutrition 12:12–26.

15. Melese K, Alemu T, Desalegn A, Urge B, Olanne A. 2025. In Vitro, Evaluation of Aflatoxin B1 Binding by Strains of Probiotic Lactic Acid Bacteria, Lactobacillus reuteri DSM 20016T and Pediococcus pentosacus DSM 20206, Isolated from Chicken Gut.

16. Tian M, Zhang G, Ding S, Jiang Y, Jiang B, Ren D, Chen P. 2022. Lactobacillus plantarum T3 as an adsorbent of aflatoxin B1 effectively mitigates the toxic effects on mice. Food Bioscience 49:101984.

17. Chen G, Fang Q, Liao Z, Xu C, Liang Z, Liu T, Zhong Q, Wang L, Fang X, Wang J. 2022. Detoxification of Aflatoxin B1 by a Potential Probiotic Bacillus amyloliquefaciens WF2020. Front Microbiol 13:891091.

18. Yan Y, Zhang X, Chen H, Huang W, Jiang H, Wang C, Xiao Z, Zhang Y, Xu J. 2022. Isolation and Aflatoxin B1-Degradation Characteristics of a Microbacterium proteolyticum B204 Strain from Bovine Faeces. Toxins 14:525.

19. Wang Y, Zhao C, Zhang D, Zhao M, Zheng D, Lyu Y, Cheng W, Guo P, Cui Z. 2017. Effective degradation of aflatoxin B1 using a novel thermophilic microbial consortium TADC7. Bioresource Technology 224:166–173.

20. Wang Y, Zhao C, Zhang D, Zhao M, Zheng D, Peng M, Cheng W, Guo P, Cui Z. 2018. Simultaneous degradation of aflatoxin B 1 and zearalenone by a microbial consortium. Toxicon 146:69–76.

21. Zhou J, Tang L, Wang J, Wang J-S. 2018. Aflatoxin B1 Disrupts Gut-Microbial Metabolisms of Short-Chain Fatty Acids, Long-Chain Fatty Acids, and Bile Acids in Male F344 Rats. Toxicological Sciences 164:453–464.

22. Mao J, Wei Y, Ni Z, Zhang J, Zhu J, Wang H. 2025. Gut microbiota-mediated bile acid transformations regulate the transport of aflatoxin B1 from the intestine to the liver in piglets. J Animal Sci Biotechnol 16:38.

23. Liu Y, Li J, Kang W, Liu S, Liu J, Shi M, Wang Y, Liu X, Chen X, Huang K. 2023. Aflatoxin B1 induces liver injury by disturbing gut microbiota-bile acid-FXR axis in mice. Food and Chemical Toxicology 176:113751.

24. Cheng, Q., Glesener, H., Sanchez Carreon, A., Voth-Gaeddert, L. E., & Krajmalnik-Brown, R. (2026). Reconsidering the Use of Dimethyl Sulfoxide for Xenobiotic-Gut Microbiota Interaction Studies. bioRxiv, 2026-08.

25. McDonald JAK, Schroeter K, Fuentes S, Heikamp-deJong I, Khursigara CM, De Vos WM, Allen-Vercoe E. 2013. Evaluation of microbial community reproducibility, stability and composition in a human distal gut chemostat model. Journal of Microbiological Methods 95:167–174.

26. Glesener H, Abdollahzadeh D, Muse C, Krajmalnik-Brown R, Weaver MA, Voth-Gaeddert LE. 2024. X-ray Irradiation Reduces Live Aspergillus flavus Viability but Not Aflatoxin B1 in Naturally Contaminated Maize.

27. Ilhan ZE, Marcus AK, Kang D-W, Rittmann BE, Krajmalnik-Brown R. 2017. pH-Mediated Microbial and Metabolic Interactions in Fecal Enrichment Cultures. mSphere 2:e00047–17.

28. Lee H-S, Salerno MB, Rittmann BE. 2008. Thermodynamic Evaluation on H_2_ Production in Glucose Fermentation. Environ Sci Technol 42:2401–2407.

29. Cheng Q, Krajmalnik-Brown R, DiBaise JK, Maldonado J, Guest MA, Todd M, Langer SL. 2023. Relationship Functioning and Gut Microbiota Composition among Older Adult Couples. IJERPH 20:5435.

30. Corbin KD, Carnero EA, Dirks B, Igudesman D, Yi F, Marcus A, Davis TL, Pratley RE, Rittmann BE, Krajmalnik-Brown R, Smith SR. 2023. Host-diet-gut microbiome interactions influence human energy balance: a randomized clinical trial. Nat Commun 14:3161.

31. Bastos-Amador P, Duarte EL, Torres J, Caldeira AT, Silva I, Salvador C, Assunção R, Alvito P, Ferreira M. 2023. Maternal dietary exposure to mycotoxin aflatoxin B1 promotes intestinal immune alterations and microbiota modifications increasing infection susceptibility in mouse offspring. Food and Chemical Toxicology 173:113596.

32. Kensler TW, Roebuck BD, Wogan GN, Groopman JD. 2011. Aflatoxin: A 50-Year Odyssey of Mechanistic and Translational Toxicology. Toxicological Sciences 120:S28–S48.

