## Supplemental Information for "Human Gut Microbiota of Healthy Individuals Show Resilience to Acute Aflatoxin B1 Exposure Despite Solvent-Driven Community Shifts"

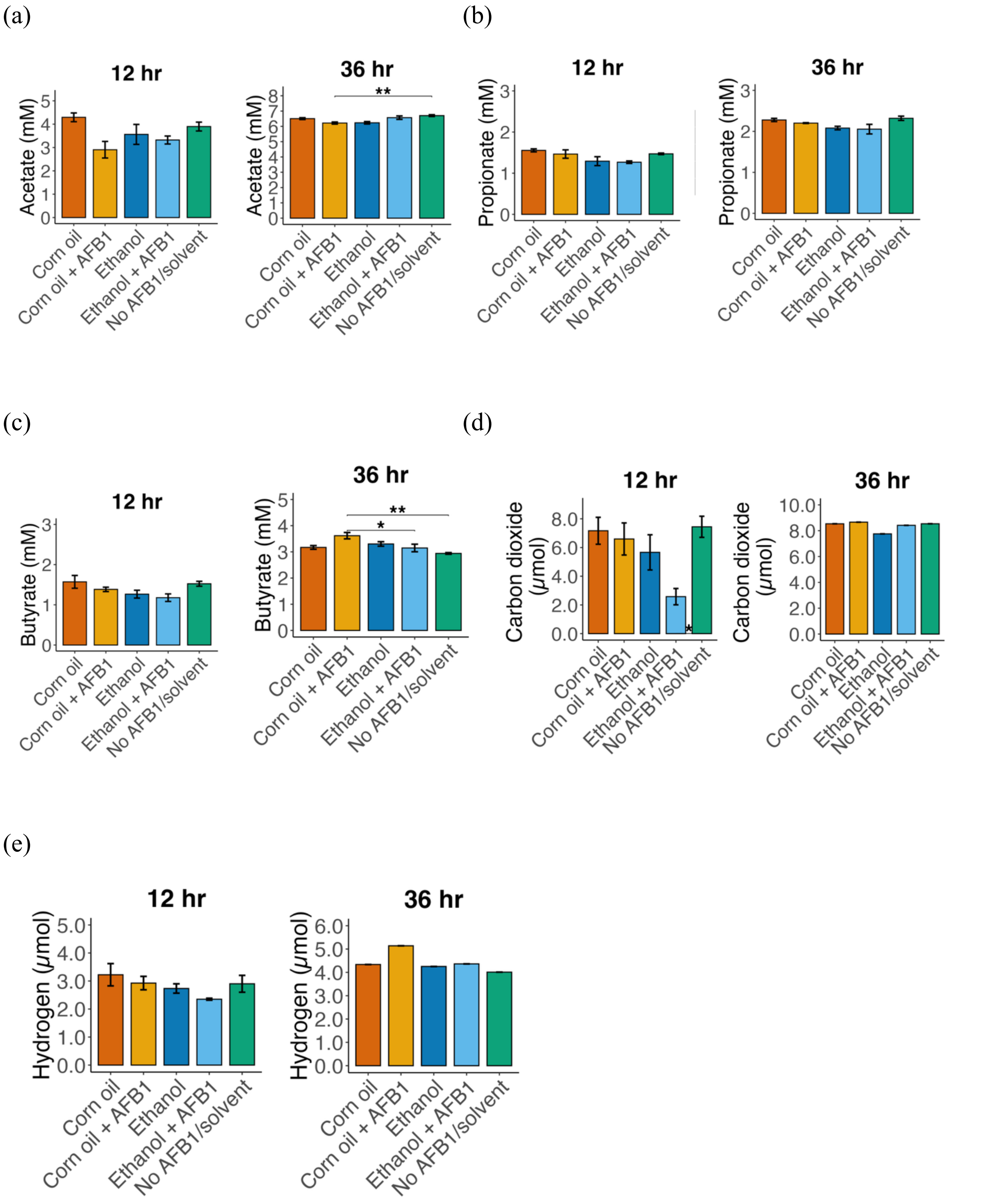


SI Figure 1. SCFA at 12 and 36 hour for the two solvent experiment. (a) acetate, (b) propionate, (c) butyrate, (d) carbon dioxide, and (e) hydrogen

(a)

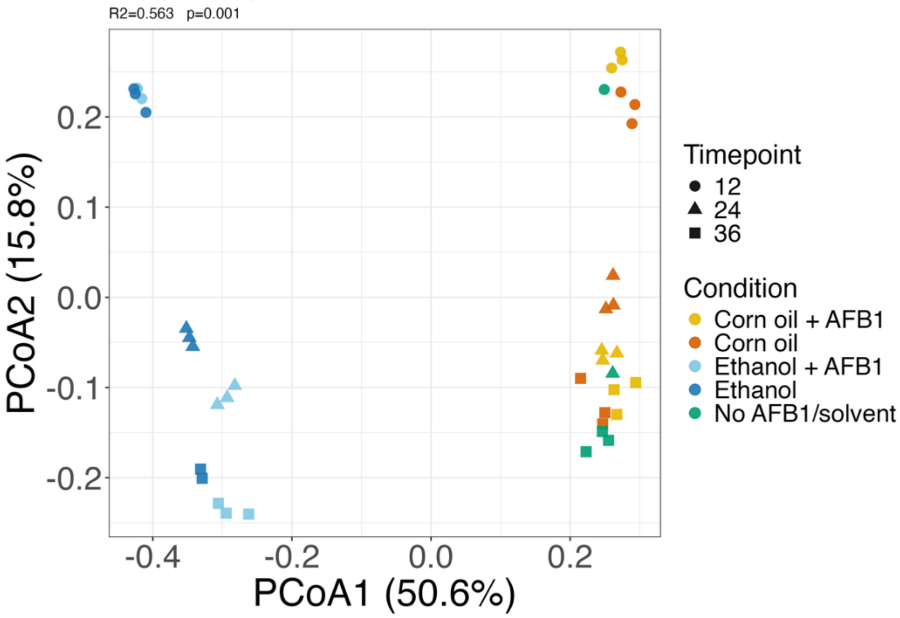


(b)

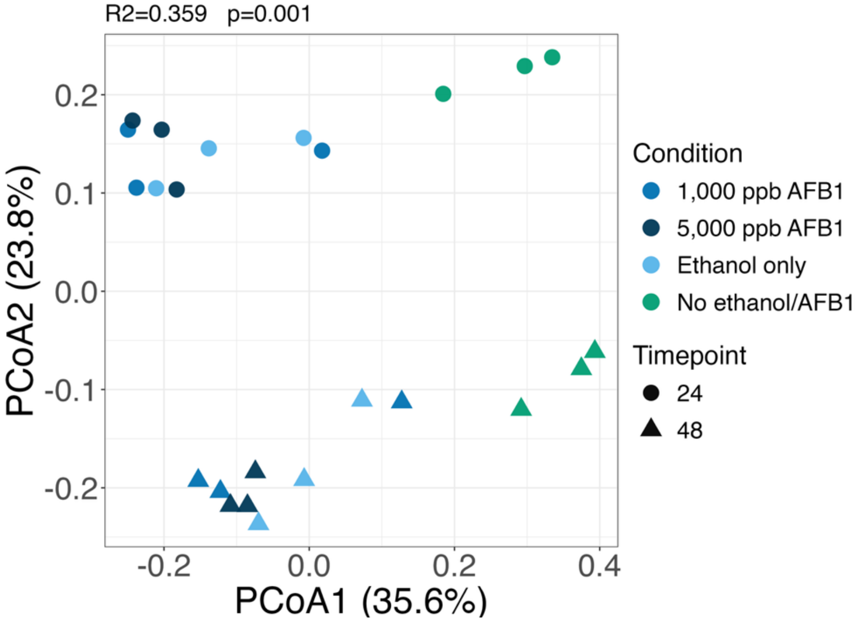


SI Figure 2. Jaccard beta diversity. (a) Solvent experiment and (b) Two-dose experiment.

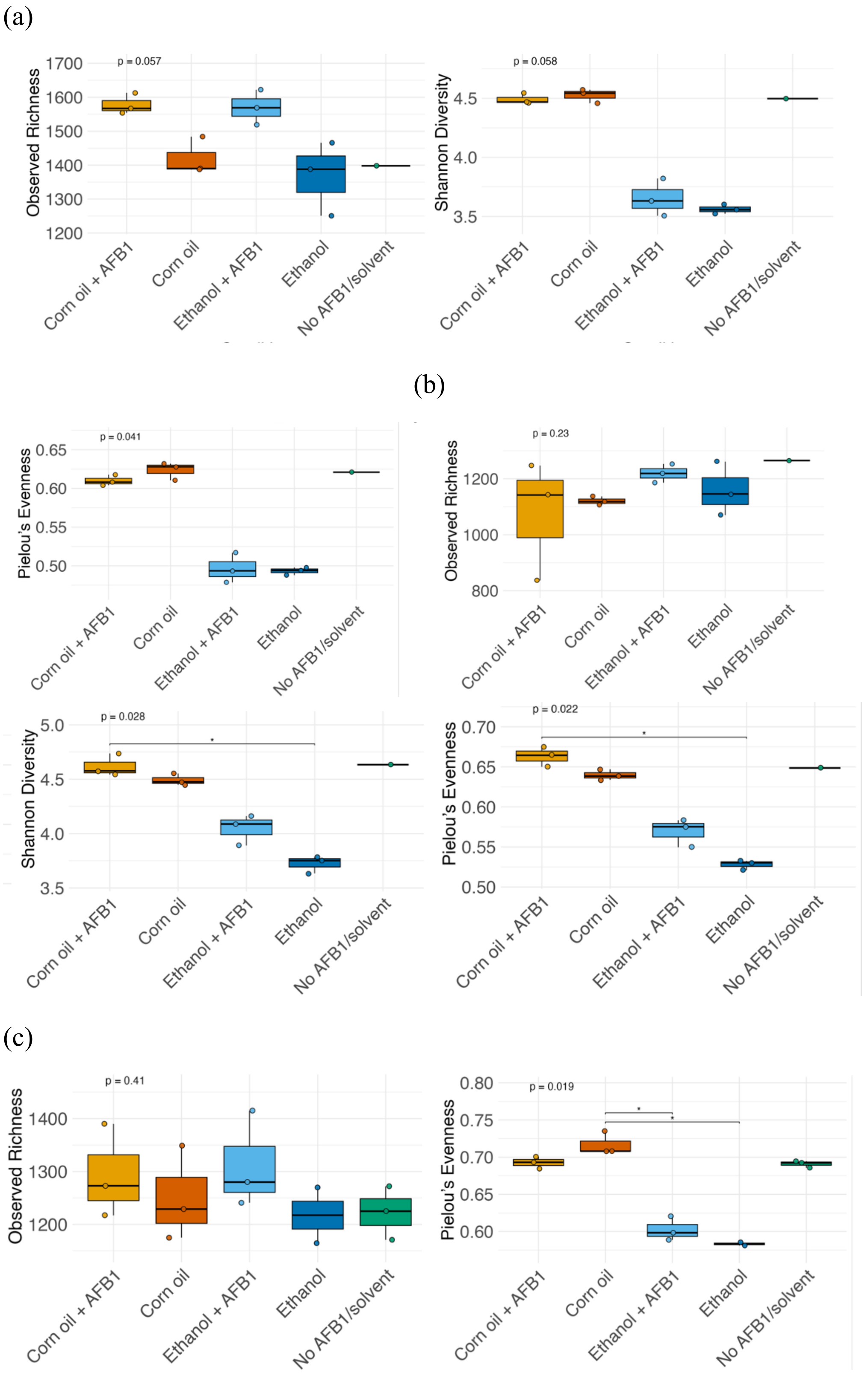


SI Figure 3. Alpha diversity metrics (Observed richness, Shannon Diversity, and Pielou's evenness) for the two solvent experiment at each timepoint (a) 12 hr (b) 24 hr and (c) 36 hr.

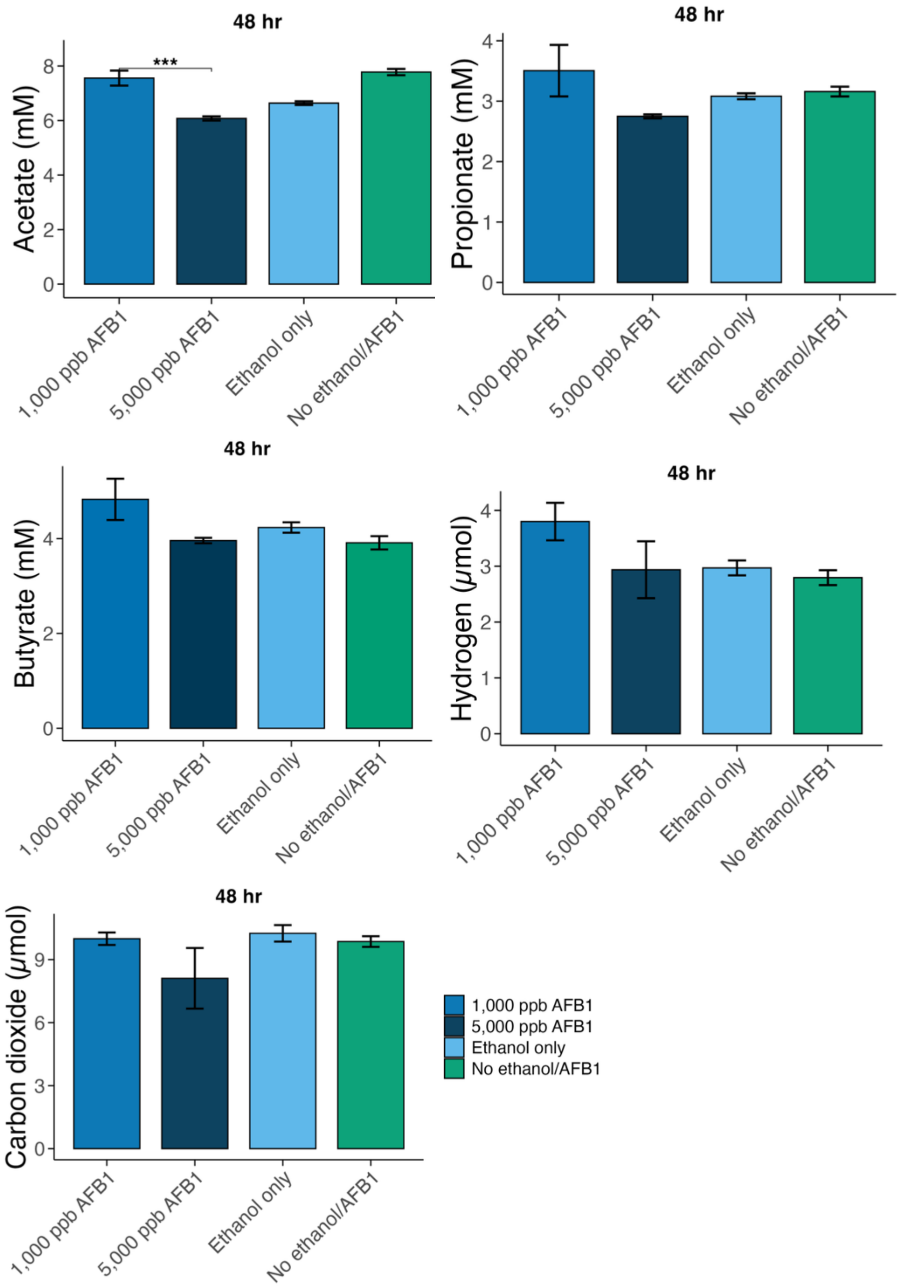


SI Figure 4. SCFA and gas at 48 hours in the two dose AFB1 experiment.
